# Morphological characterization of moulting in the Atlantic horseshoe crab *Limulus polyphemus* and evolutionary insights into moulting patterns in Arthropoda

**DOI:** 10.64898/2026.02.27.708456

**Authors:** Kenneth Kim, Sinéad Lynch, Harriet B. Drage, Jonathan Antcliffe, Ariel D. Chipman, Allison C. Daley, Marc Robinson-Rechavi

## Abstract

Arthropods must periodically moult their exoskeleton to permit growth, a conserved developmental process whose morphological and behavioural execution varies widely among lineages. Horseshoe crabs (Limulidae) are members of Xiphosura, a chelicerate lineage with a fossil record extending as far back as the Ordovician and provide a valuable comparative framework for studying the evolution of moulting strategies in Arthropoda. Despite their importance, detailed morphological characterization of moulting in horseshoe crabs remains scarce, limiting developmental studies and broader comparative analyses. Here, we provide a detailed morphological characterization of the moulting process in the Atlantic horseshoe crab *Limulus polyphemus*. Morphological changes in specific anatomical structures, including the anterior margin of the prosoma, lateral spines and dorsal spinous process of the opisthosoma, were observed during the moulting process. By tracking these morphological markers, such as retraction of the epidermis from the cuticle and degree of corrugation of the epidermis, we were able to identify individuals in the early and late pre-moult stage, predict the onset of ecdysis, and distinguish post-moult and intermoult stages. We compare ecdysis patterns in *L. polyphemus* with other arthropod taxa, both extant and fossil. We find that, despite differences in behavioural execution, ecdysis in *L. polyphemus* shares features with other chelicerates, and that both phylogenetic signal and convergent patterns are evident across Arthropoda. This study offers a robust, non-invasive method for determining moult stages in juvenile horseshoe crabs and provides insights into the diversity and constraints of ecdysis in Arthropoda.

## Introduction

Arthropods, including marine chelicerates such as horseshoe crabs, must periodically moult their exoskeleton in order to grow. Moulting is orchestrated by tightly regulated hormonal cues, such as sesquiterpenoids and ecdysteroids, that trigger a cascade of physiological, biochemical, and morphological events culminating in the exit of the animal from its old exoskeleton, or ecdysis (Campli et al., 2024; Cheong et al., 2015; Daley and Drage, 2016; Qu et al., 2018; Truman and Riddiford, 2019). Ecdysteroids coordinate the sequence of pre-moult, ecdysis, and post-moult events, including cuticle degradation, mineral resorption, and formation of the new cuticle (Mykles, 2011; Mykles and Chang, 2020). While moulting is a conserved developmental process across Arthropoda, the morphological and behavioural execution of ecdysis is strikingly diverse. For example, the positioning of ecdysis suture lines (Snodgrass, 1947), the directionality of emergence from the old cuticle, and preparatory behaviours before ecdysis vary across groups (Sullivan et al., 2024). Understanding these differences is essential for reconstructing how moulting has been shaped by ecological pressures and body plan architectures over evolutionary time and provides valuable insights into the evolution of developmental strategies in Arthropoda.

Horseshoe crabs (Limulidae) are members of Xiphosura, a chelicerate lineage with a rich fossil record extending back to the Ordovician (Rudkin et al., 2008; Van Roy et al., 2015), and are often regarded as “living fossils” owing to their striking morphological conservatism (Castellano et al., 2025; Kin and Błażejowski, 2014) although this characterization has been increasingly questioned (Bicknell, 2019; Bicknell et al., 2019; Bicknell and Pates, 2020). Nevertheless, xiphosurans provide crucial comparative context for inferring ancestral arthropod character states and for examining broader evolutionary patterns across Arthropoda, given the placement of Chelicerata as the sister group to Mandibulata (Regier et al., 2010). Previous work on horseshoe crabs has documented moulting primarily in the contexts of conservation and aquaculture (Chatterji et al., 2004; Mishra, 2009; Schreibman and Zarnoch, 2009; Tanacredi et al., 2009), growth and recovery from injury (Bicknell and Cuomo, 2024), and post-embryonic development with an emphasis on the occurrence and timing of ecdysis (Sekiguchi, 1988; Sekiguchi et al., 1988). However, morphological characterization of its moult cycle, encompassing the full sequence of pre-moult, ecdysis, and post-moult events, remains poor compared to other groups.

In well-studied arthropod groups such as decapod crustaceans, decades of research have established detailed moult staging systems based on morphological criteria (Corteel et al., 2012; Drach and Tchernigovtzeff, 1967; Gorissen and Sandeman, 2022; Kim et al., 2024). Among these, Drach’s classification (Drach and Tchernigovtzeff, 1967) has become the gold standard for decapod crustaceans, providing clear nomenclature and sub-stage definitions that facilitate comparative studies. In contrast, the moulting process in horseshoe crabs lacks a standardized set of morphological criteria for moult staging. Previous studies of the Atlantic horseshoe crab, *Limulus polyphemus*, have relied on indirect proxies such as the number of days since the last moult and body size increments to infer moult stage (Sekiguchi, 1988; Sekiguchi et al., 1988) or external descriptions such as based on coloration (Winget and Herman, 1979). These approaches can be confounded by individual variability and environmental influences in the development of *L. polyphemus* (Haug and Rötzer, 2018), particularly when the instar stage is unknown and thus potentially prone to under sampling or oversampling of the moulting stage. This issue is particularly pronounced given that the relative duration of specific moult stages can vary substantially between species (Gorissen and Sandeman, 2022; Kim et al., 2024; Volovych et al., 2025). Moult stage characterizations are particularly important in comparative analyses of moulting, as the genes regulating moulting are expressed in a hierarchical cascade (Ashburner, 1974; Campli et al., 2024; Truman and Riddiford, 2019). Because developmental processes are continuous, the use of arbitrary temporal sampling points might not necessarily correspond to biologically meaningful stages, particularly when stage durations vary across species (Levin et al., 2012; Roux et al., 2015). These differences are not simply proportional to, for example, moult duration. The absence of moult stage criteria not only limits developmental and physiological studies within horseshoe crabs but also hampers our ability to make meaningful cross-species comparisons of moulting processes across Arthropoda. A comprehensive morphological characterization of the moult cycle is therefore essential both for species-specific research and for broader evolutionary analyses, enabling the identification of conserved and divergent patterns in moulting across arthropod lineages.

To address this gap, we first characterized the moulting process in *Limulus polyphemus* through detailed examination of changes in specific anatomical structures such as the anterior margin of the prosoma, the dorsal spinous process of the opisthosoma, and the lateral spines we were able to reliably distinguish different moult stages and their progression. These include epidermal retraction from the cuticle (apolysis), the degree of epidermal corrugation, tanning and hardening of the new cuticle, and the formation of the sutural gape during ecdysis. Together, these features allowed us to distinguish early and late pre-moult, predict the onset of ecdysis, and identify post-moult and intermoult individuals. We also documented behavioural observations associated with the onset and progression of ecdysis and compared ecdysis patterns of *L. polyphemus* across other arthropod taxa. Our findings provide a precise, non-invasive method for determining the moult stages in juvenile *L. polyphemus*. We then compared ecdysis in *L. polyphemus* with other taxa, characterising both shared features with other chelicerates despite differences in behavioural execution, i.e. phylogenetic signal, and convergent patterns among arthropods with similar head shields.

## Materials and methods

Juvenile horseshoe crabs used in this study were bought from private suppliers and reared in an aquarium research laboratory of the Institute of Earth Sciences, University of Lausanne. Tanks containing the horseshoe crabs were maintained with a water temperature of 21 °C and salinity of 25.1 psu (⁓25.1 ppt or a specific gravity of 1.019). Total ammonia-N were kept below 0.5 mg L-1, nitrite-N below 0.15 mg L-1, and nitrates as close to zero as possible. All tanks received artificial sunlight for 8 h per day. They were fed once daily with commercial pellet feed (JBL Pronovo, Neuhofen, Germany).

Moult stages of the horseshoe crab were differentiated and characterised by observing the morphological changes at the anterior margin of the prosoma (ventral side). Additional structures were also observed such as the dorsal spinous processes of the opisthosoma, and lateral spines (Figure 1). A total of 10 individuals were observed measuring 1.5 – 18 cm in length from the anterior margin of the prosoma to the posterior tip of the telson with an average increase of 3.2 cm in length after moulting with varying numbers of observations per stage. As moulting was not synchronised among individuals, data collection was constrained by the stages present during the duration of observation. For some individuals, all moult stages were documented, whereas for others, only a single stage could be observed, as larger individuals required considerably longer time to reach the next moult. The number of individuals observed per stage are presented in each results section. Each individual was tagged by using cyanoacrylate adhesives and labels were placed on the side of the carapace (Figure 1). Suitability and safety of the adhesives were based on results from mussels (Harrtman et al., 2016). Individuals were observed using an Olympus SZX10 stereomicroscope with an observed magnification of 0.63–6.3× and were imaged using an SC50 5-megapixel colour 118 camera (Olympus Life Science Solutions, Olympus, Tokyo, Japan) with Preciv Image Analysis Software (version 1.2; Evident Corporation, Tokyo, Japan). Ecdysis was also characterized by the timing of the formation of a sutural gape and accompanying behavioural observations were recorded.

**Figure 1.**
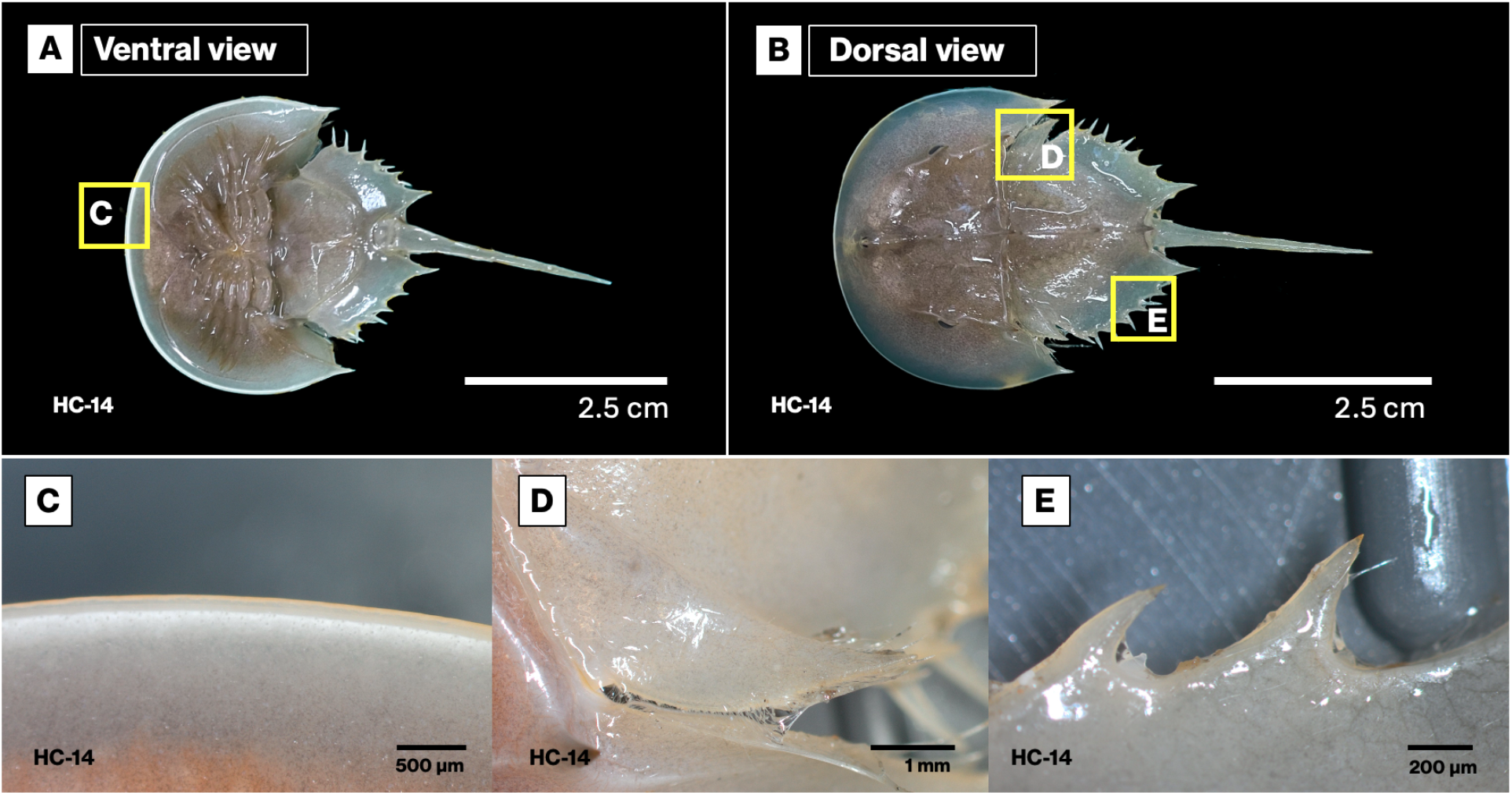
Overview of morphological markers used in this study on the (A) ventral and (B) dorsal side of the horseshoe crab (HC). Microscope images of the morphological markers of *L. polyphemus* during intermoult: (C) anterior margin of the prosoma, (D) dorsal spinous process, and (E) lateral spines. Individual identifiers (HC-n) are shown at the bottom left of each photo.

We also compared ecdysis patterns in *L. polyphemus* with those reported across other arthropod lineages by summarising key moulting traits. These traits include the direction of egress during ecdysis (i.e. the direction in which the animal exits the exuvia) and the position of the suture gape or ecdysial suture, which refers to the location at which the exuvia splits open during moulting (Box 1). Trait information was retrieved from moultDB (https://www.moultdb.org) and published literature for various taxa across several arthropod lineages (Andrew and Patankar, 2010; Drage et al., 2023b; Faasch, 1967; Gaban and Farley, 2002; García-Bellido and Collins, 2004; Haupt, 1982; Iwasaki et al., 2000; Kim et al., 2024; Minch, 1977; Moysiuk and Caron, 2024; Norton and Kethley, 1994; Phlippen et al., 2000; Price and Holdich, 1980; Rieder, 1972; Snodgrass, 1947; Tetlie et al., 2008) (Figure 6). Ecdysial suture position was categorized based on the location and orientation of the primary opening through which the animal exits the exoskeleton.

### Box 1

Definition of terms for egress direction and ecdysial suture position.

#### Egress direction

##### Anterior

The animal exits the exuvia by moving forward along the anterior–posterior axis, with the head and anterior body regions emerging first through an anterior opening.

##### Posterior

The animal exits by moving backward relative to the exuvia, with posterior body regions emerging first.

##### Dorsal

Emergence occurs upward relative to the body axis, with the dorsal side of the body lifting and moving out from the exuvia.

##### Anterior–dorsal

The animal emerges through a combined forward and upward movement, with the anterior body regions exiting first while also lifting dorsally. This represents an oblique trajectory rather than a purely anterior or dorsal direction.

##### Biphasic

Egress occurs in two sequential phases. Typically, one body region emerges first, followed by a second phase in which the remainder of the body exits. This is observed in Isopods where the animal sheds the posterior (rear) half of their exoskeleton before the anterior (front) half.

#### Ecdysial suture position

##### Anterior

The ecdysial opening is located at the anterior margin of the body, often involving the head region or anterior cephalic shield. Egress occurs forward through a frontal split.

##### Lateral

Lateral opening refers to involvement of the lateral margins of the body in the ecdysial opening, either through propagation of an initial rupture or as part of the primary opening. The opening usually originates along a dorsal or anterior axis and subsequently extends laterally during emergence.

##### Posterolateral

The ecdysial opening is located along the posterolateral margin of the cephalothorax, forming a split at the posterior edge of the carapace. In brachyuran crustaceans, this position corresponds to the posterior boundary of the cephalothorax, which appears posterior relative to other decapods due to reduction and ventral folding of the pleon.

##### Transverse

The ecdysial opening runs across the body perpendicular to the anterior–posterior axis. In contrast to a complete circumferential or full transverse split, this suture may be partial without fully separating the body into anterior and posterior halves.

##### Circumferential

The suture encircles the body partially or completely, forming a ring-like opening that allows the animal to emerge by separating dorsal and ventral cuticle layers.

##### Mid-transverse

A transverse suture located approximately at the mid-body region separating anterior and posterior regions of the body as often observed in Isopods.

##### Epicranial to mid-dorsal

A longitudinal suture extending from the head region (epicranium) along the dorsal midline of the body.

##### Dorsal head–trunk junction

The ecdysial opening is located dorsally at the boundary between the head and trunk regions.

##### Anterior dorsolateral suture

The ecdysial opening is located along the dorsolateral margin of the head, typically near the boundary between the head and the first trunk segment, forming a hinge-like opening during ecdysis. In some taxa such as in diplopods the position of this suture may be shifted posteriorly and involve the second trunk tergite rather than the head itself.

##### Suture of cephalothorax

The opening occurs along the dorsal region of the cephalothorax (fused head and thorax), typically forming a carapace split that allows the animal to emerge anteriorly or dorsally.

It should be noted that the overview in Box 1 is not exhaustive but provides a comparative summary of ecdysis traits across representative arthropod taxa. Moulting trait information was mapped onto a simplified arthropod phylogeny based on Legg et al. (2013), to visualise their distribution across major lineages. We highlight lineages that share with horseshoe crabs a domed, pronounced and horseshoe-shaped cephalic shield with curved or convex anterior and lateral margins, which in harpid and trinucleid trilobites are further expanded into a wide, flattened cephalic brim or fringe.

## Results

### Intermoult

During the intermoult stage, the most noticeable feature is the progressive tanning of the cuticle, reflected by increased sclerotization and changes in pigmentation across the three morphological markers (*n* = 5) (Figure 1C–E). Compared to the post-moult stage, the cuticle becomes pigmented and rigid, losing its pale and wrinkled appearance. The cuticle is characterized by having an off-white to a light brown tone across the prosoma, dorsal spinous process, and lateral spines (Figure 1). The onset of intermoult is thus defined by a rigid cuticle and browner appearance across all regions. The duration of intermoult until pre-moult is approximately 60–120 days with shorter duration in smaller individuals than in larger ones.

### Early pre-moult

Early pre-moult is characterized by the appearance of an ecdysial gap (*n* = 6) (Figure 2, red arrows). This marks the separation of the epidermis from the old cuticle, a process called apolysis. The gap first appears along the anterior margin of the prosoma and is already prominent, whereas in the dorsal spinous process and lateral spines it appears more distinctly as an emerging ecdysial line (Figure 2B, C). It should be noted that in larger individuals, the cuticle is considerably thicker and more opaque, making the early stages of pre-moult more difficult to detect until the gap has widened sufficiently. This gap continues to expand as the epidermis becomes increasingly corrugated toward the later stages of pre-moult. Estimated duration of early pre-moult ranges from 18 to 23 days.

**Figure 2.**
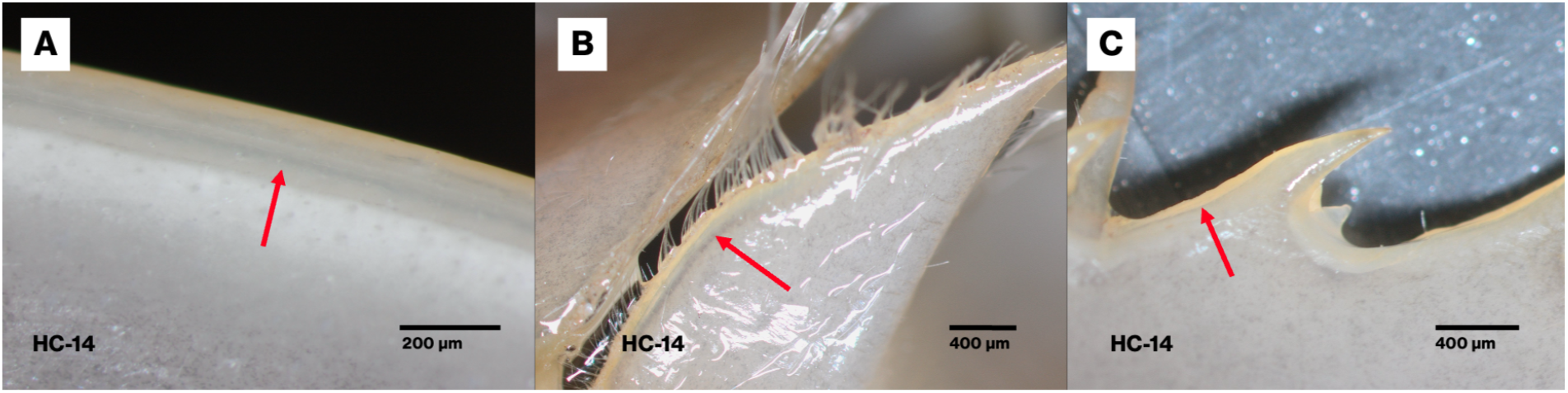
Microscope images of the morphological markers of *L. polyphemus* during early pre-moult. (A) Anterior margin of the prosoma, (B) dorsal spinous process and (C) lateral spines. Red arrows indicate the ecdysial gap. Individual identifiers (HC-n) are shown at the bottom left of each photo.

### Late pre-moult

Late pre-moult is characterized by the presence of epidermal corrugations and the secretion of the new cuticle, which will continue to thicken (*n* = 9) (Figure 3A: red arrow). Moreover, the retraction of the epidermis is more evident, forming a space which is visible on the spinous process and the lateral spines (red arrows in Figure 3B and C). The outline of the separating epidermis in the lateral spines also exhibits a whiter appearance. The presence of the epidermal corrugations and the wider ecdysial gap seen in the lateral spines is a good indicator that the animal will moult in approximately 1-2 days with smaller individuals having shorter duration than larger ones.

**Figure 3.**
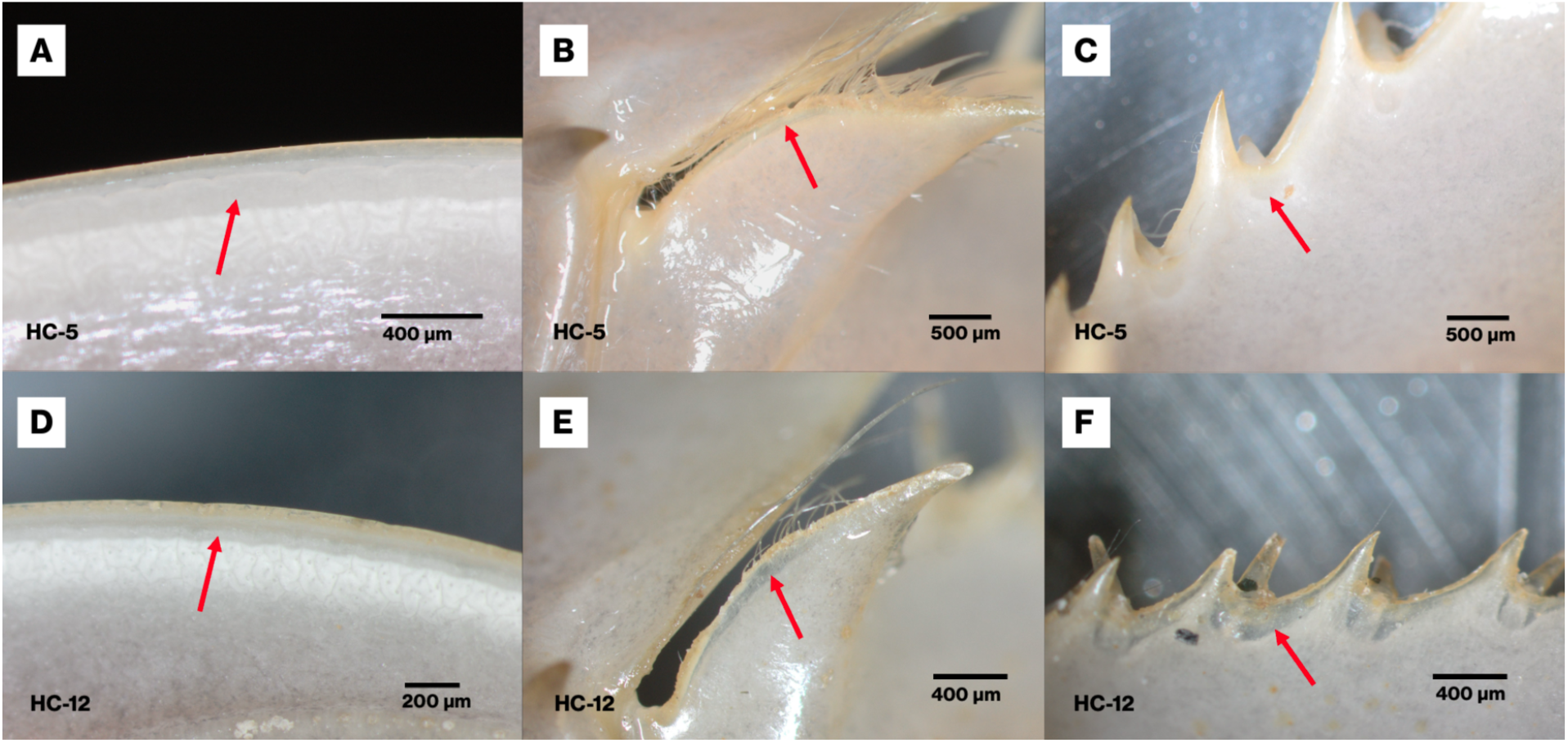
Microscope images of the morphological markers of *L. polyphemus* during late pre-moult. (A,D) Anterior margin of the prosoma, (B,E) dorsal spinous process and (C,F) lateral spines. Top panel A–C: HC-5 (length = 6 cm), bottom panel D–F: HC-12 (length = 3.8 cm). Red arrows indicate the ecdysial gap. Individual identifiers (HC-n) are shown at the bottom left of each photo.

### Ecdysis

Prior to ecdysis, the animal typically remains on the substrate surface and exhibits a digging posture characterized by the anterior portion of the body being angled downward, with the head shield directed toward the substrate. During this period, the animal exhibits a digging behaviour while it uses its telson for stabilization ([Video 1](https://zenodo.org/records/20185971/files/Video_1_small_individual.mp4?download=1)). During ecdysis, the animal undergoes repeated flexion of the body at the prosoma–opisthosoma articulation, producing a characteristic wiggling motion ([Video 2](https://zenodo.org/records/20185971/files/Video_2_small_individual_ecdysis.mp4?download=1)). Simultaneously, visible contractions and expansions of the soft carapace suggest the generation of internal hydrostatic pressure. Together, these movements appear to facilitate the gradual emergence of the animal from the exuvia. This behaviour is observed in smaller individuals (approximately 2–5 cm in length). In contrast, larger individuals (12–17 cm) display a different pattern of activity, often moving across the substrate rather than remaining stationary in a fixed position ([Video 3](https://zenodo.org/records/20185971/files/Video_3_large_individual.mp4?download=1)). Ecdysis is initiated by the formation of a sutural gape at the anterior margin of the carapace (Figure 4A), which progressively enlarges as the animal expands and exerts pressure to rupture the exuviae. The rupturing of the sutural gape can initially be asymmetric. In Figure 4B the animal has ruptured most of the left side but not the right side of the sutural gape. In individuals measuring approximately 15 cm, we estimate the total duration of the ecdysis process within 8-12 hours, from the initial appearance of the sutural gape to complete emergence.

**Figure 4.**
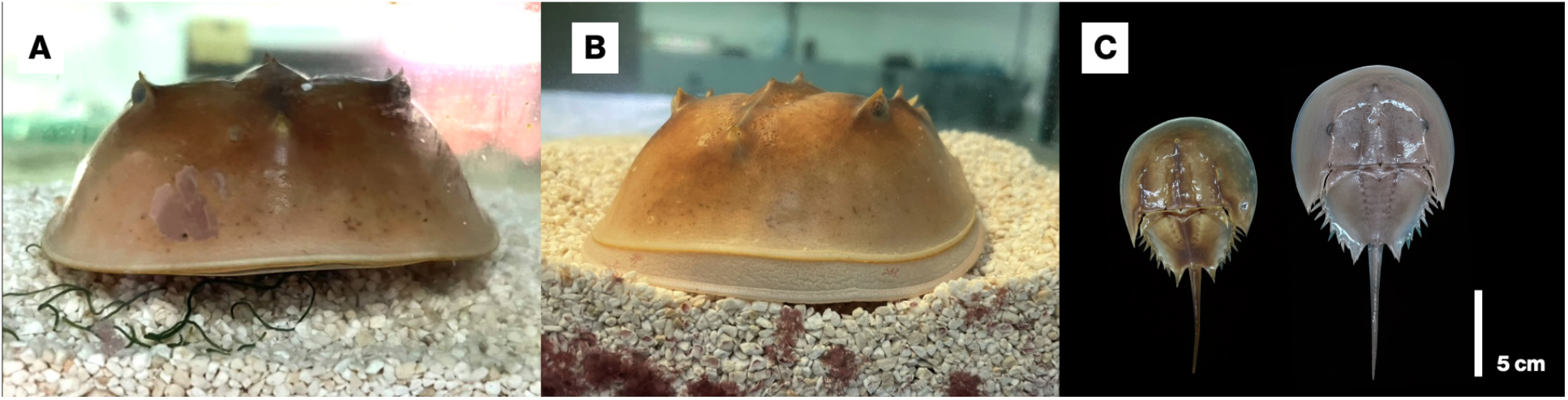
Images of *L. polyphemus* during ecdysis. (A) Opening sutural gape at the anterior end of the prosoma, (B) ecdysis, and (C) old exuvia (left) and animal after ecdysis (right).

### Post-moult

Immediately after ecdysis, the cuticle of *Limulus polyphemus* is thin, wrinkled, and pale, which is particularly noticeable at the anterior margin of the carapace (*n* = 5) (Figure 5A). The spinous process and lateral spines also appear lighter and paler compared to pre-moult (Figure 5). The tips of the spines are already darker and harder than the surrounding carapace, suggesting that sclerotization and tanning occur more rapidly in these regions than elsewhere on the exoskeleton. During post-moult, coloration gradually becomes light brown over several days, beginning as a pale cuticle that darkens first at the anterior end of the prosoma and progresses posteriorly toward the lateral spines. For this observation, we used the coloration of the dorsal spinous process and lateral spines as the point of reference in the duration of post-moult until intermoult which is estimated from 3–7 days with smaller individuals having shorter duration than larger ones.

**Figure 5.**
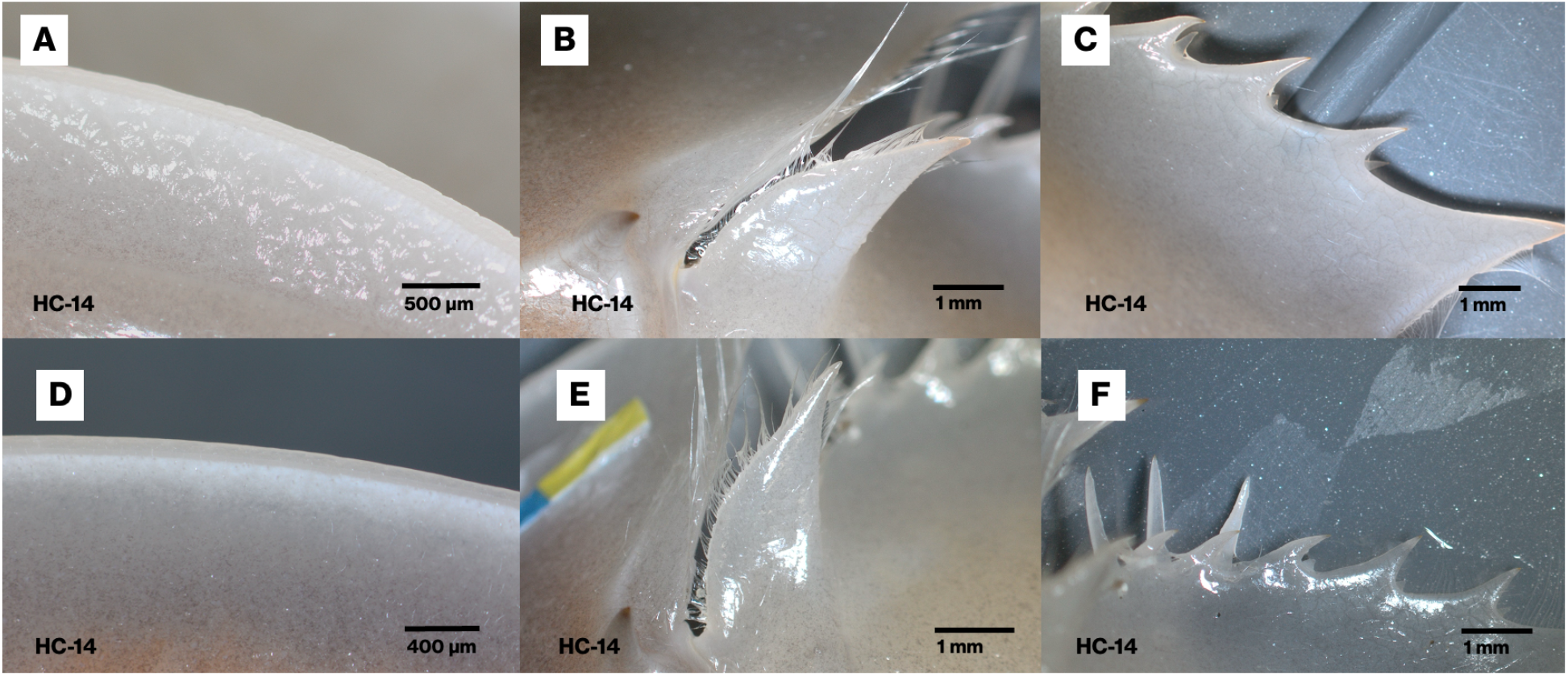
Microscope images of the morphological markers of *L. polyphemus* during postmoult. (A,D) Anterior margin of the prosoma, (B,E) dorsal spinous process and (C,F) lateral spines. Top panel (A–C) 24 hrs after moulting; bottom panel (D–F) 48 hrs after moulting. Individual identifiers (HC-n) are shown at the bottom left of each photo.

### Comparison of ecdysis patterns between *Xiphosurida* and other Arthropods

The overview obtained by comparing egress direction and location of moulting suture between Arthropoda clades highlights the diversity of ecdysis patterns present across the phylum and reveals shared and distinct ecdysis traits among extant and extinct taxa. Taxa that share a domed, pronounced and horseshoe-shaped cephalic shield, all have moulting sutures located anteriorly on the cephalon, facilitating egress in the anterior direction. This is the case for both extant and extinct taxa that are distantly related on the phylogenetic tree, including the stem lineage clade Radiodonta, Trilobita, the chelicerate clade Xiphosura, and the pancrustacean clade Branchiopoda. Insects and myriapods have highly canalised moulting suture and egress directions, with the former being characterised by epicranial to mid-dorsal sutures facilitating anterior-dorsal egress, and the latter favouring dorsolateral head sutures and anterior egress. In contrast, moulting traits are highly variable for crustaceans. Chelicerates also show some degree of variation in moulting traits, but with more of a predominance for anterior sutures and egress direction.

## Discussion

In this study, we provide a detailed morphological characterisation of the moult cycle in the horseshoe crab *Limulus polyphemus*. Although morphological characterisations of moulting are lacking for xiphosurids, aspects of the moulting process have been investigated in other chelicerates, albeit with different emphases. Previous studies have primarily characterised the behavioural sequence and mechanics of ecdysis, describing changes in posture, carapace splitting, and the order of appendage extraction during emergence, as documented for scorpions and spiders (Gaban and Farley, 2002; Minch, 1977). Other work has focused on cellular and ultrastructural processes associated with cuticle renewal during apolysis, particularly in relation to sensilla and hair regeneration in spiders and solifuges (Harris, 1977; Haupt, 1982). Comparable ultrastructural descriptions of the moulting process are also available for mites, where apolysis, early epicuticular deposition in discrete plaques that subsequently fuse into a continuous layer, epidermal folding, and the determination of cuticular ridges have been documented in detail (Crowe, 1975; Mothes-Wagner, 1984). Consistent with these observations, we found that the formation of epidermal corrugations and pre-ecdysial cuticle secretion in *L. polyphemus* closely resembles processes described in other arthropods, including the hemimetabolous insect *Blattella germanica* (Cruz et al., 2007), the branchiopod *Triops cancriformis* (Rieder, 1972), and the isopod *Oniscus asellus* (Price and Holdich, 1980). However, in *L. polyphemus*, formation of the new cuticle becomes apparent only during late premoult, when epidermal corrugations are visible, and whether earlier phases of epicuticular deposition, as observed in mites, have occurred is not resolved at the current resolution used for this study.

Across arthropods, apolysis and secretion of the new cuticle prior to ecdysis are recurring features, and convolutions or corrugations of the newly forming cuticle are evident before emergence. During moulting, apolysis marks the separation of the epidermis from the old cuticle and the formation of an apolytic space (Locke, 2001; Locke and Huie, 1979). The epidermis then begins secreting the initial layers of the new cuticle, while moulting fluid released into the apolytic space digests the inner layers of the old cuticle, allowing resorption and recycling of chitin and protein components (Merzendorfer and Zimoch, 2003; Moussian, 2010). Throughout this process, the newly synthesised cuticle is protected from degradation by moulting enzymes through the action of cuticle-associated proteins such as Knickkopf (Chaudhari et al., 2011). Because the new cuticle must ultimately accommodate a greater surface area, it is initially laid down in a folded configuration within the confined exuvial space. Following ecdysis, the cuticle remains wrinkled, pale and soft during the post-moult period and gradually expands and hardens. Together, these observations indicate that epidermal folding and cuticular corrugation represent a conserved biomechanical response for increased surface area during cuticle renewal, indicating that the moult stages described here may extend to other xiphosurids and possibly to chelicerates more broadly.

We also observed that ecdysial behaviour differs between smaller and larger individuals, which likely reflects size-dependent morphological constraints associated with body architecture. This pattern is consistent with previous findings on post-embryonic allometric changes in *Limulus polyphemus*, where shifts in prosomal and thoracetron development across moult stages are hypothesized to be linked to transitions in locomotor behaviour and life mode, from primarily surface interaction to more active benthic burrowing and foraging (Bicknell and Cuomo, 2025). As body size increases, changes in cuticle thickness, body mass, and overall geometry are expected to alter mechanical demands during ecdysis, potentially necessitating different behavioural strategies to facilitate emergence from the old exoskeleton. In addition, external hydrodynamic forces may also contribute to size-dependent differences in ecdysial behaviour. Aquatic organisms are known to adopt postures that minimise drag or generate negative lift to remain attached under turbulent flow conditions (Martinez, 2001; Maude and Williams, 1983; Pates and Drage, 2024; Weissenberger et al., 1991). Experimental and computational fluid dynamics studies on *L. polyphemus* demonstrate that carapace morphology and orientation strongly influence lift and drag forces under flowing water to maintain contact with the substrate (Davis et al., 2019). Ontogenetic changes in size and shape may therefore alter how individuals experience hydrodynamic forces during ecdysis, particularly in benthic or shallow-water environments. While it remains unclear whether differences in ecdysial behaviour are driven primarily by shifts in life mode, hydrodynamic influences, or their interaction, these observations suggest that growth-related allometry plays an important role in shaping the behavioural strategies associated with moulting in *L. polyphemus*.

Across Arthropoda, moulting strategies are diverse, and ecdysis patterns reflect both phylogenetic and convergent signals (Figure 6). Within Chelicerata, including horseshoe crabs, scorpions, and spiders, ecdysial sutures or suture gapes occur at the anterior and lateral margins of the prosoma despite differences in the egress direction and behavioural execution of ecdysis. Although the comparative summary presented here is not exhaustive, suture gape in the anterior and lateral margin appears to represent a recurrent feature among arachnids and xiphosurids with clear prosoma–opisthosoma tagmosis. A similar ecdysis pattern is also inferred to represent the ancestral condition in mites, based on outgroup comparison. It is widespread across several acariform lineages, including endeostigmatic mites, eupodine Prostigmata, and Oribatida, even though these taxa lack the pronounced external tagmosis observed in non-mite arachnids (Northon and Kethley, 1994). In contrast, derived mite groups within Acariformes exhibit alternative ecdysial sutures, including transverse and circumferential suture gapes associated with posterior emergence, indicating secondary diversification of moulting strategies within the clade (Faasch, 1967; Rocket and Woodring, 1972; Northon and Kethley, 1994). Whether the anterior prosomal suture gape reflects an ancestral feature across Chelicerata remains to be clarified by future work on other chelicerates especially those with divergent body plans, such as pycnogonids, and opilionids (Chipman, 2025; Dunlop and Lamsdell, 2017).

**Figure 6.**
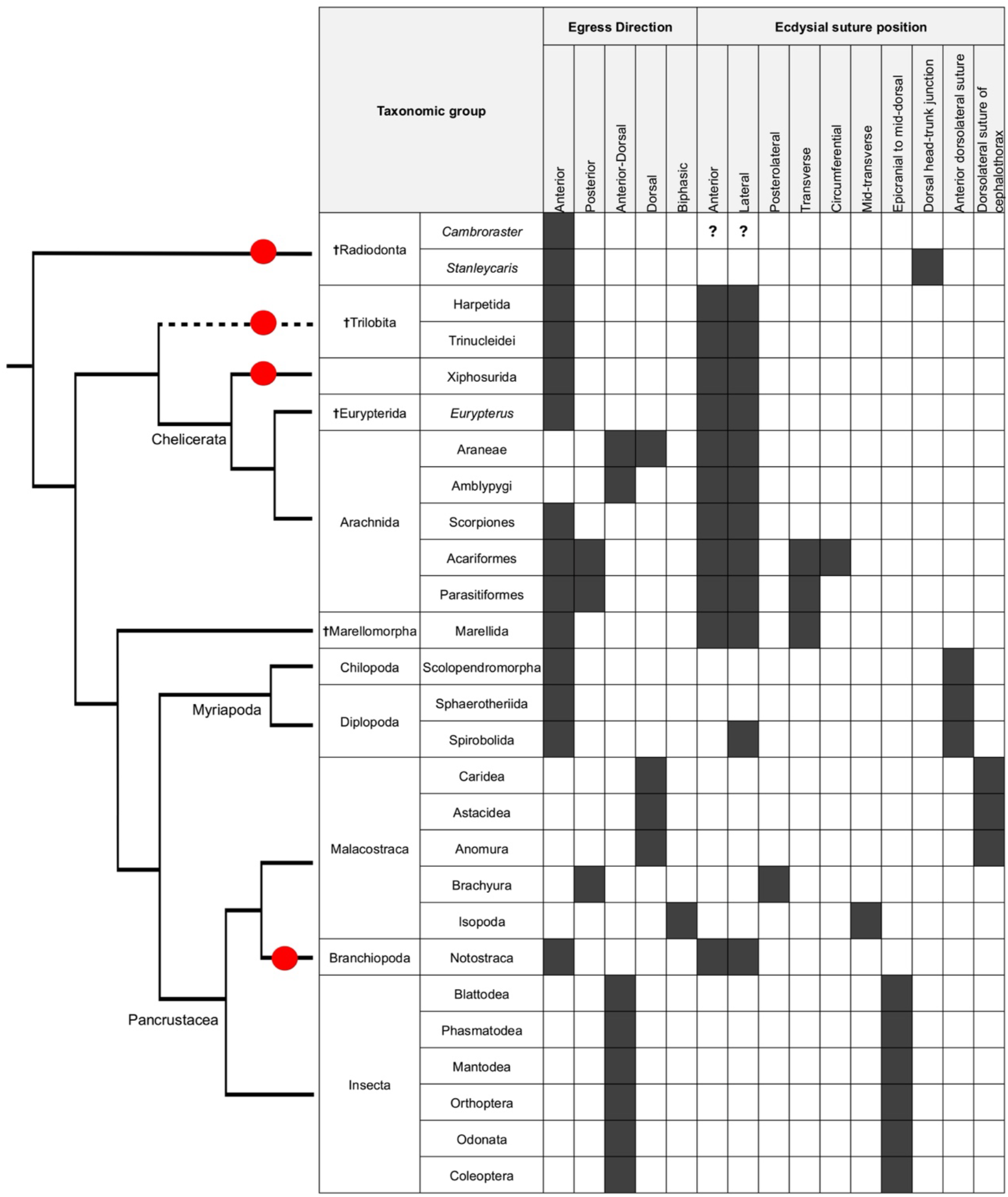
Phylogenetic distribution of egress direction during moulting and ecdysial suture position across major arthropod lineages. The simplified phylogeny (left) is adapted from Legg *et al*. 2013 and illustrates relationships among major extinct (**†**) and extant groups, with red disks indicating occurrence of taxa characterised by a domed and pronounced horseshoe-shaped head shield. The uncertain phylogenetic position of trilobita is indicated by a broken line. The accompanying matrix (right) shows moulting traits for representative taxa. Filled cells indicate documented occurrences of a given trait, whereas empty cells indicate absence or lack of available data. Question mark denotes uncertain moulting traits in extinct taxa. Definition of terms are described in methods Box 1.

Beyond Chelicerata, similar ecdysial sutures can arise convergently. Ecdysis in *Limulus polyphemus*, or specifically emergence through an anterior marginal suture in the headshield, is comparable to that described in *Triops*, a branchiopod crustacean with a similar domed and horseshoe-shaped cephalic shield. The appearance of anterior marginal sutures in *Triops* indicates that similar ecdysial sutures can evolve independently in non-chelicerate taxa in response to mechanical demands imposed by comparable exoskeletal architecture. Similar moulting patterns also been inferred for several extinct arthropod lineages that possess rigid, expanded headshields with anterior marginal sutures, including harpetid and trinucleid trilobites, as well as other less diverse trilobite clades (Daley and Drage, 2016; Drage et al., 2019; Du et al., 2024) which is hypothesized to minimize structural compromise during ecdysis (Esteve et al., 2021). The repeated evolution of horseshoe-shaped and shield-like cephalic morphologies across phylogenetically distant arthropods, such as in the Cambrian stem arthropod *Cambroraster*, trilobites, xiphosurids, and notostracans, suggests convergent responses to similar ecological adaptations (Liu et al., 2020). However, the precise ecological functions of these expanded headshields remain unresolved. These morphologies have been variously interpreted as adaptations for benthic or nektobenthic lifestyles (Botton and Jr, 2003; Fortey, 2014), plowing (Bergström, 1972), burrowing (Eldredge, 1970), sediment interaction, or hydrodynamic stability (Davis et al., 2019), suspension feeding (Fortey and Owens, 1999) and these hypotheses continue to be actively investigated (Pates and Drage, 2024). Moreover, quantitative analyses of trilobite moulting demonstrate that moulting behaviour in this group was highly variable and cannot be reliably predicted from cephalic shape, facial suture type, or overall body proportions, even within single species (Drage, 2024; Drage et al., 2023). These results indicate that, in trilobites, anterior exuviation was one viable strategy among several rather than a fixed outcome of headshield morphology, which indicates a more plastic moulting strategy. In contrast, extant arthropod lineages with broadly similar cephalic morphology, such as xiphosurids and notostracans, exhibit more stereotyped moulting strategies. This suggests that while anterior exuviation may represent a recurrent solution to the mechanical challenges imposed by rigid and expanded headshields, the degree to which this solution becomes canalized is different between trilobites and extant taxa, potentially reflecting different post-ecdysial modification strategies, lineage specific cuticular structures, or tighter coupling between morphology and moulting behaviour.

Taken together, these findings point towards a broader evolutionary pattern in which ecological pressures associated with particular life modes favoured changes in exoskeletal architecture, such as the evolution of domed and horseshoe-shaped head regions. Such morphological changes would, in turn, impose mechanical constraints on ecdysis, necessitating modifications to suture placement and moulting behaviour. Conversely, moulting is not merely constrained by morphology but may itself influence exoskeletal structure by restricting the range of morphological configurations compatible with repeated cuticle renewal. Feedback between ecology, biomechanics, and developmental processes may therefore reinforce particular morphological solutions over evolutionary time (Brandt, 2002). Ecologically advantageous traits that conflict with the mechanics of ecdysis may require developmental decoupling from the moult cycle, as exemplified by holometabolous insects, in which membranous structures such as wings which lack epidermal cells are produced only after a terminal moult (Belles, 2019; Truman, 2019). Expanded morphological and behavioural annotation of moulting across the arthropod tree of life will be essential to further test this hypothesis and to examine the evolution, diversification, and constraint of moulting in Arthropoda.

## Author contributions

KK designed the study and experiment, led formal analysis and data collection, figure preparation, and manuscript writing (original draft, review, and editing). JA was responsible for aquarium maintenance and animal husbandry and assisted with experimental procedures. All authors reviewed the draft manuscript. SL, HD, ADC, ACD, and MRR reviewed, edited and revised the original draft and the final manuscript. All authors read and approved the submitted manuscript.

## Acknowledgments

The authors would like to thank the members of the Arthropod Moulting Evolution Project: Robert Waterhouse, Giulia Campli, Michele Leone, Sagane Joye-Dind, Valentine Rech de Laval, Olga Volovych, Idan Sheizaf, and Asya Novikova, for their invaluable support throughout this study. We are grateful to Farid Saleh, Pierre Gueriau, Gaetan Potin and Nora Corthésy for their assistance in the aquarium lab. We would also like to thank Emelyne Gaudichau for sharing her observations on the moulting of stick insects. This work was supported by funding from the Swiss National Science Foundation (SNSF) Sinergia grant [grant number 198691].

## Supplementary information

Video files in this study are available at Zenodo: 10.5281/zenodo.18402874

